# Role of intraluteal and intrauterine prostaglandin signaling in LH-induced luteolysis in pregnant rats

**DOI:** 10.1101/2022.05.09.490003

**Authors:** Akshi Vashistha, Medhamurthy Rudraiah

## Abstract

Luteal dysfunctions lead to fertility disorders and pregnancy complications. Normal luteal function is regulated by many factors, including luteinizing hormone (LH). The luteotropic roles of LH have been widely investigated but its role in the process of luteolysis has received little attention. LH has been shown to have luteolytic effects during pregnancy in rats. Stocco et al. have demonstrated the role of intraluteal prostaglandins (PGs) in LH-mediated luteolysis. However, the status of PG signaling in the uterus during LH-mediated luteolysis remains unexplored. In this study, we have examined the effect of LH-mediated luteolysis on luteal and uterine PG synthesis machinery and genes associated with activated luteal PGF_2α_ signalling and uterine activation during different stages (mid and late) of pregnancy. Further, we analysed the effect of overall PG synthesis machinery blockage on LH-mediated luteolysis during late-pregnancy. Unlike the mid-stage of pregnancy, the expression of genes involved in PG synthesis and responsivity in late-stage pregnant rats’ luteal and uterine tissue increase post repeated administration of LH. Since the cAMP/PKA pathway mediates LH-mediated luteolysis, we analyzed the effect of inhibition of endogenous PG synthesis on the cAMP/PKA/CREB pathway, followed by the analysis of the expression of markers of luteolysis. Inhibition of endogenous PG synthesis did not affect the cAMP/PKA/CREB pathway. However, in the absence of endogenous PGs, luteolysis could not be activated to the full extent. Our results suggest that endogenous PGs may contribute to LH-mediated luteolysis, but this dependency on endogenous PGs is pregnancy stage dependent. These findings advance our understanding of the molecular pathways that regulate luteolysis.

## Introduction

In female mammals, the process of luteolysis is complex and varies from species to species. Luteinizing hormone (LH) induces luteolysis in rats during the mid and late-stage of pregnancy (Vashistha *et al*., 2021). Stocco and colleagues emphasized the role of intraluteal prostaglandin (PG) F_2α_ in LH-induced luteolysis (Stocco and Deis, 1998). However, they did not analyse the role of uterine PGF_2α_ signaling in LH-mediated luteolysis. In this study, we have analysed the contribution of endogenous PG arising from both corpus luteum (CL) and the uterus in LH-mediated luteolysis.

PGF_2α_ is a well-established luteolysin and plays a crucial role during luteolysis in many species, including bovines (Metcalf *et al*., 1992), primates (Stouffer *et al*., 2013) and rodents (Bjurulf *et al*., 1998). In rodents, PGF_2α_ of both the luteal and uterine origin are involved in luteolysis (McCracken *et al*., 1999). The uterus remains in a quiescent state throughout pregnancy until the beginning of parturition (Renthal *et al*., 2010). Uterine activation in rats is associated with increased expression of a set of genes encoding prostaglandin synthesis machinery and contraction-associated proteins (CAPs) (JRG *et al*., 2000; Mitchell *et al*., 2005). The prostaglandin synthesis machinery comprises of four main enzymes including cyclooxygenase (*Cox*) 1, *Cox2*, PGF synthase (*Pgfs*), and carbonyl reductase 1 (*Cbr1*). PGs are synthesized through the hydrolysis of membrane phospholipid into arachidonic acid by the action of phospholipases (Ricciotti and FitzGerald, 2011). Arachidonic acid is further converted to PGH2 by the enzyme COX1 and COX2, this step is the rate limiting step in the production of PGs (Ricciotti and FitzGerald, 2011). PGH2 is quite unstable in nature and is further converted into different PGs namely PGE2, PGD2, PGF2, PGI2 by PGE2 synthase, PGD2 synthase, PGFS and PGI2 synthase, respectively (Ricciotti and FitzGerald, 2011). Cox1 and Cox2 are bifunctional enzymes that contain both cyclooxygenase and peroxidase activity. The two cyclooxygenase isoforms, COX1 and COX2, are targets of nonsteroidal anti-inflammatory drugs (NSAIDs) (Allaj *et al*., 2013). NSAIDs binds and inactivate the COX, which results in a decrease in the peripheral and central prostaglandin production (Allaj *et al*., 2013). Many NSAID have been used for this purpose including, indomethacin, naproxen, diclofenac sodium (DIC), ketoprofen, aspirin and many others (Allaj *et al*., 2013). DIC [2-(2,5-dichlorophenyl) amino] benzene acetic acid, is one of the most widely used NSAID for treatment of pain and inflammation (van Walsem *et al*., 2015). Its specificity for both the isoforms of cyclooxygenase is almost the same and is metabolized in the liver (Tomic *et al*., 2008). It has been used for reproductive studies to analyse the effects of suppression of prostaglandins on oxytocin sensitivity and parturition (Chan, 1983).

Some of the CAPs known to be up-regulated at the time of uterine activation include oxytocin (Lefebvre *et al*., 1992), fibronectin (Tuo and Bazer, 1996), interleukin-1 (IL-1), osteopontin (OPN) (Waterhouse *et al*., 1992), P-glycoprotein (Axiotis *et al*., 1991), and insulin-like growth factor binding protein 2 (IGFBP-2) (Rutanen, 2000). Cytokines also play an important role in uterine contractility, cervical ripening, and birth (Orsi and Tribe, 2008). Reports suggest an association of nuclear factor kappa-light-chain-enhancer of activated B cells (NF-κB) and IL-1β with the contraction of the uterus (Orsi and Tribe, 2008). Other cytokines like transforming growth factor-β1 (TGF-β1), IL-1α, interferon-γ (IFN-γ) and tumor necrosis factor-α (TNF-α) have also been implicated in uterine activation (Orsi and Tribe, 2008).

For sufficient activation of the uterus along with activation of CAP, repression of the relaxation system is required (Challis *et al*., 2000). Inducers of myometrial relaxation include P_4_, NO, PTH-related peptide, CRH, and others (Challis *et al*., 2000). A study in 2003 (Girotti and Zingg, 2003) utilized microarray technique to identify different factors that may regulate the process of uterine relaxation. Among the expression of other genes identified in the analysis, a gene named *Ugn*, which codes uroguanylin (UGN) protein, decreased its expression up to 20-fold immediately before parturition (Jaleel *et al*., 2002). UGN ligand binds to its transmembrane enzyme-linked receptor guanylate cyclase C (GC-C) present on the uterus (Jaleel *et al*., 2002). Upon ligand binding, GC-C catalyses the conversion of GTP to cGMP, and cGMP relaxes smooth muscle cells by inhibiting Ca^2+^ entry (Carvajal *et al*., 2000). P_4_ has been shown to up-regulate the expression of EP2 to maintain uterine quiescence, however expression of EP2 and EP4 becomes down regulated at the time of parturition in rodents to potentiate the process of contraction (Breyer *et al*., 2001).

We conducted experiments to determine the involvement of PGF_2α_ signaling during LH-mediated luteolysis. Our specific objectives were to analyse: (1) the effect of repeated administration of exogenous LH on PG synthesis and its signaling in the CL and uterus during mid and late-pregnancy in rats and (2) the effect of inhibition of endogenous PG synthesis on LH-mediated luteolysis and PGF_2α_ signaling in the CL and uterus.

## Material and Methods

### Reagents

Ovine LH and Cetrorelix (Cet) were kind gifts from Professor M R Sairam (University of Montreal, Quebec, Canada) and Asta Medica (Frankfurt, Germany), respectively. Trizol (93289) and Diclofenac (D6899) were purchased from Sigma Aldrich Co. (Bangalore, India). P_4_ RIA kit (IM1188) was procured from Beckman Coulter (CA, USA). Antibody against cyclic adenosine monophosphate (cAMP), CV-27 pool was obtained from Dr. A.F. Parlow, NHPP (CA, USA). PKA assay kit (ADI-EKS-390A) was purchased from Enzo Life Sciences (NY, USA). Table 1 contains details of the antibodies employed. All the other reagents were purchased from Sigma Aldrich Co. (Bangalore, India) or sourced from local distributors.

**Table 1.** List of antibodies used for immunoblotting

| Si. No. | Antibody | Company | Catalogue number | Dilution | Secondary antibody |
| --- | --- | --- | --- | --- | --- |
| 1. | GAPDH D16H11 | CST | #5174 | 1:1000 | Anti-rabbit |
| 2. | Beclin 1 | CST | #3738 | 1:1000 | Anti-rabbit |
| 3. | LC3I/II D3U4C0 | CST | #12741 | 1:1000 | Anti-rabbit |
| 4. | Cleaved caspase 3 | CST | #9661 | 1:1000 | Anti-rabbit |
| 5. | pCREB Ser133 | CST | #9198 | 1:1000 | Anti-rabbit |
| 6. | CREB 48H2 | CST | #9197 | 1:1000 | Anti-rabbit |

### Animals

The Institutional Animal Ethics Committee approved experimental protocol involving rats. Rattus norvegicus (Harlan-Wistar strain) were housed in a controlled environment and kept under 12 h light and 12 h dark cycles with ad libitum access to food and water. To obtain pregnant rats, 2-3 month old female virgin rats were cohabitated with male rats. Vaginal smears were screened daily for the presence of sperm. We designated the day of the appearance of sperm as DOP 1. After establishing pregnancy, female rats were housed individually for further experimentation. The gestation period in our colony of rats is 23 days. We collected blood and CL at suitable intervals depending on the experiment.

### LH-mediated induction of luteolysis in pregnant rats

We carried out studies in the late (DOP 19) and mid-stage (DOP 8-10) pregnant rats. Previously published protocol was followed (Vashistha *et al*., 2021). Briefly, late-stage pregnant rats were assigned into two groups with three animals/group. On DOP 19, each group received either four i.p. injections of Veh (100 μL of 1× PBS) or LH (10 μg dissolved in 100 μL of 1× PBS) at 08:00, 09:00, 10:00 and 11:00 h. On DOP 20, rats from both groups received one i.p. injection of LH (10 μg). Circulating LH levels are high during mid-pregnancy, and CL functions are dependent on LH during this period (Loewit *et al*., 1969). To rule out the participation of endogenous LH during exogenous administration of LH, Cet, a GnRH receptor antagonist, was administered to inhibit endogenous levels of LH. The tested dose and duration of Cet treatment were reported previously (John *et al*., 2016). Administration of 150 μg/Kg BW of Cet every 12 h on DOP 8-9 of pregnancy resulted in luteolysis and pregnancy loss (John *et al*., 2016). We assigned rats on DOP 8 to four different groups with three rats/group. Rats in the first and second groups received four s.c. injections of either Veh (100 μL of 1× PBS) or Cet (150 μg/Kg BW) at 12 h intervals for 48 h beginning on DOP 8, respectively. Post Cet treatment, we administered one i.p. injection of Veh on DOP 10. Rats in the third and fourth groups received Cet treatment as per the second group. Furthermore, on DOP 10, rats from the third and fourth group received one or four injections of LH (10 μg: at hourly intervals), respectively. The first, second, third, and fourth groups are referred to as Veh, Cet+Veh, Cet+LH, and Cet+4LH groups, respectively. Blood and CL were collected from rats during both stages, 40 min after the last Veh/LH injection.

### Effect of PGF_2α_ synthesis blockade on LH induced luteolysis

Endogenous PGs were inhibited by employing DIC during late-pregnancy to confirm the role of PG signalling during LH-induced luteolysis. Previously published protocol with few modifications (Chan, 1983) was followed. Briefly, we assigned 18 days old pregnant rats (late-pregnant stage) into two groups (3 animals/group). Rats in both groups received DIC by oral gavage three times daily [6 am (1 mg), 12 pm (0.5 mg), and 6 pm (0.5 mg)] on days 18-19 of pregnancy. On day 20 of pregnancy, both the groups received one injection of DIC (6 am) and a single i.p. injection of LH (10 μg: dissolved in 100 μl of 1X PBS) at 11 am. Additionally, rats in the second group received four injections of LH (10 μg: dissolved in 100 μl of 1X PBS) at hourly intervals on day 19 of pregnancy as well. Blood, CL, and uterine tissue were collected from rats during both stages, 40 min after the last Veh/LH injection.

### Tissue and blood processing

Fat was cleared from the CL and uterine tissue upon collection. The tissues were then flash-frozen in liquid nitrogen and stored at -70° C for RNA and protein analysis. Blood was further processed into plasma and serum and stored at -20° C until analysis.

### qPCR

Total RNA from CL tissues was isolated using TRI reagent as per the manufacturer’s recommendations. The qPCR analysis was performed as described previously (Vashistha *et al*., 2021). Table 2 contains the list of primers used in the qPCR analysis.

**Table 2.**
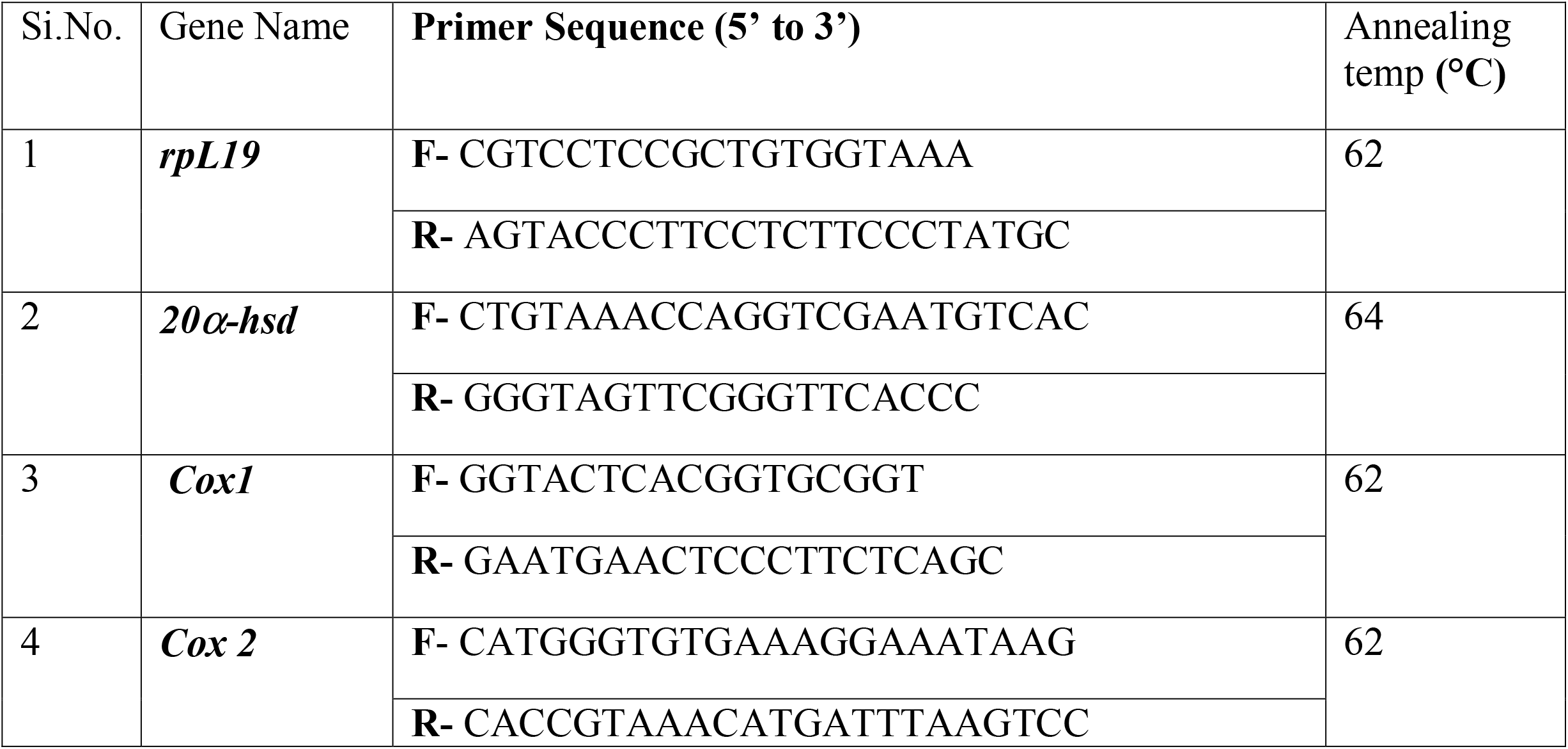
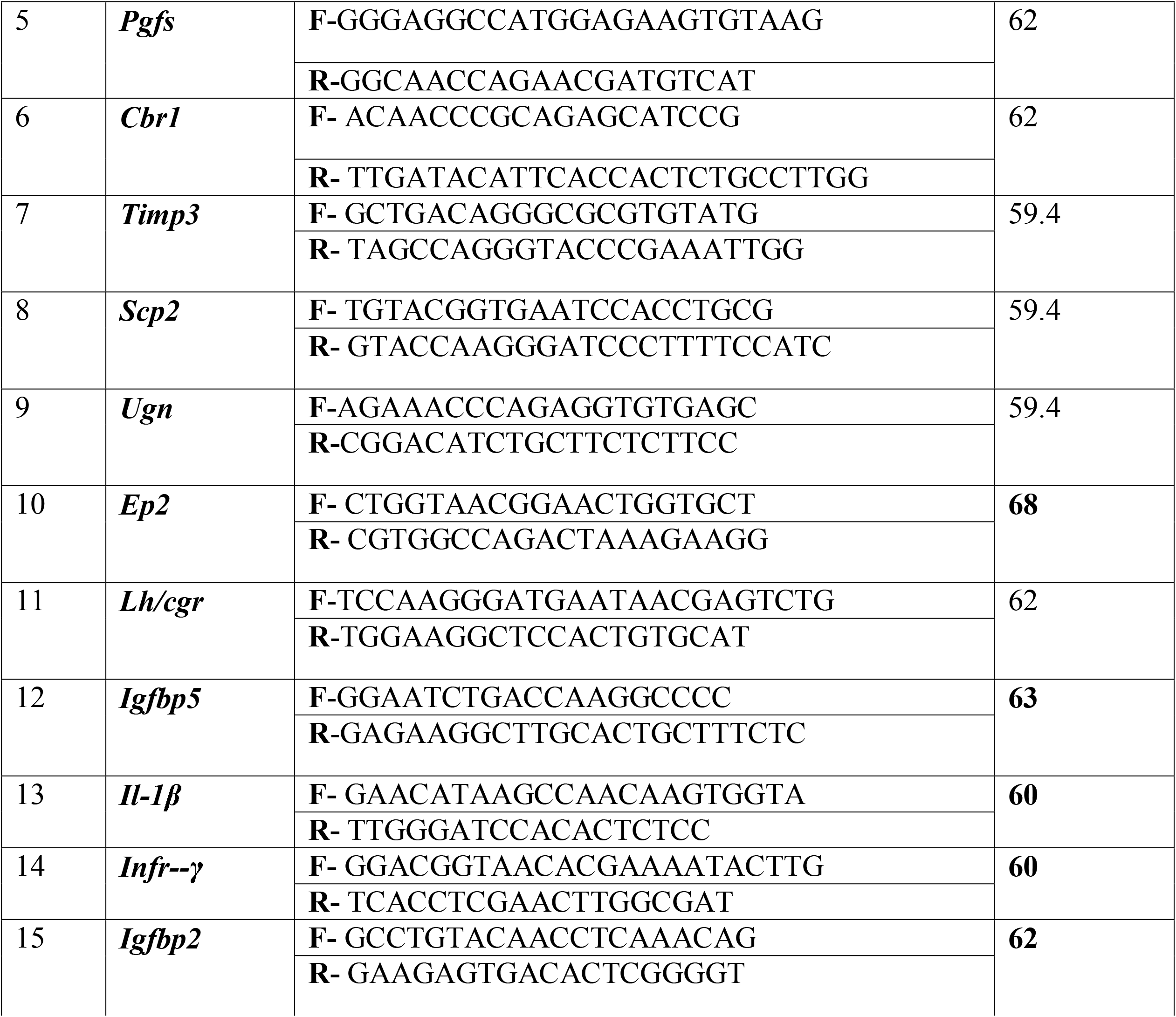
List of primers used for real-time PCR

### Immunoblotting

Immunoblot analysis of the total protein lysates from CL tissue was carried out as per the procedures reported previously (Kunal *et al*., 2012). Table 1 contains the list of antibodies used in the immunoblotting analysis.

### RIA of cAMP

cAMP lysate preparation and the assay protocol followed have been described previously (Kunal *et al*., 2012). The antibody (CV-27) at a dilution of 1:30,000 (∼ 40% binding) was used for the assay of samples.

### P_4_ assay

Serum P_4_ levels were determined by employing a commercially available RIA kit as per the manufacturer’s instructions. The inter-and intra-assay coefficients of variation were found to be ≤ 9%.

### Statistical analysis

All data is expressed as mean±SEM. The graphs were plotted and analysed using GraphPad Prism® 5 software (GraphPad Software, Inc., La Jolla, CA, USA). A two-tail paired ‘t’ test with 95% confidence intervals was used for statistical analysis of the results between two groups. One-way analysis of variance (ANOVA) and Bonferroni post-tests with 95% confidence intervals were used to compare three or more groups. A p-value < 0.05 was considered to be statistically significant.

## Results

### Effect of repeated LH administration on the expression of PGF_2α_ biosynthesis machinery during pregnancy

We examined the participation of endogenous PGF_2α_ signaling in LH-induced luteolysis. The expression of genes involved in PGF_2α_ synthesis (*Cox1, Cox2, Pgfs* and *Cbr1)* was analyzed at the transcript level in the CL and uterine tissue. During late pregnancy, repeated administration of LH decreased the luteal expression of *Cox1* (p<0.01) and *Cox2* (non-significant) [responsible for the synthesis of precursor prostaglandin, PGH2] (Fig. 1A, B). However, the expression of genes *Pgfs* (p<0.01) and *Cbr1* (p<0.05) increased in the CL, the enzymes coded by the genes are responsible for the conversion of PGH2 and PGE2 to PGF_2α,_ respectively (Fig. 1C, D). Transcript levels of PGF_2α_ biosynthesis genes increased [*Cox1* (non-significant), *Cox2 (p<0*.*01), Pgfs (p<0*.*02), and Cbr1* (non-significant)] in the intact uterus post LH treatment (Fig. 1 E-H) during late pregnancy. No change occurred in the levels of PGF_2α_ biosynthesis genes in the CL (Fig. 1I-L) and uterine tissue collected from mid-pregnant rats (data not shown).

**Figure 1.**
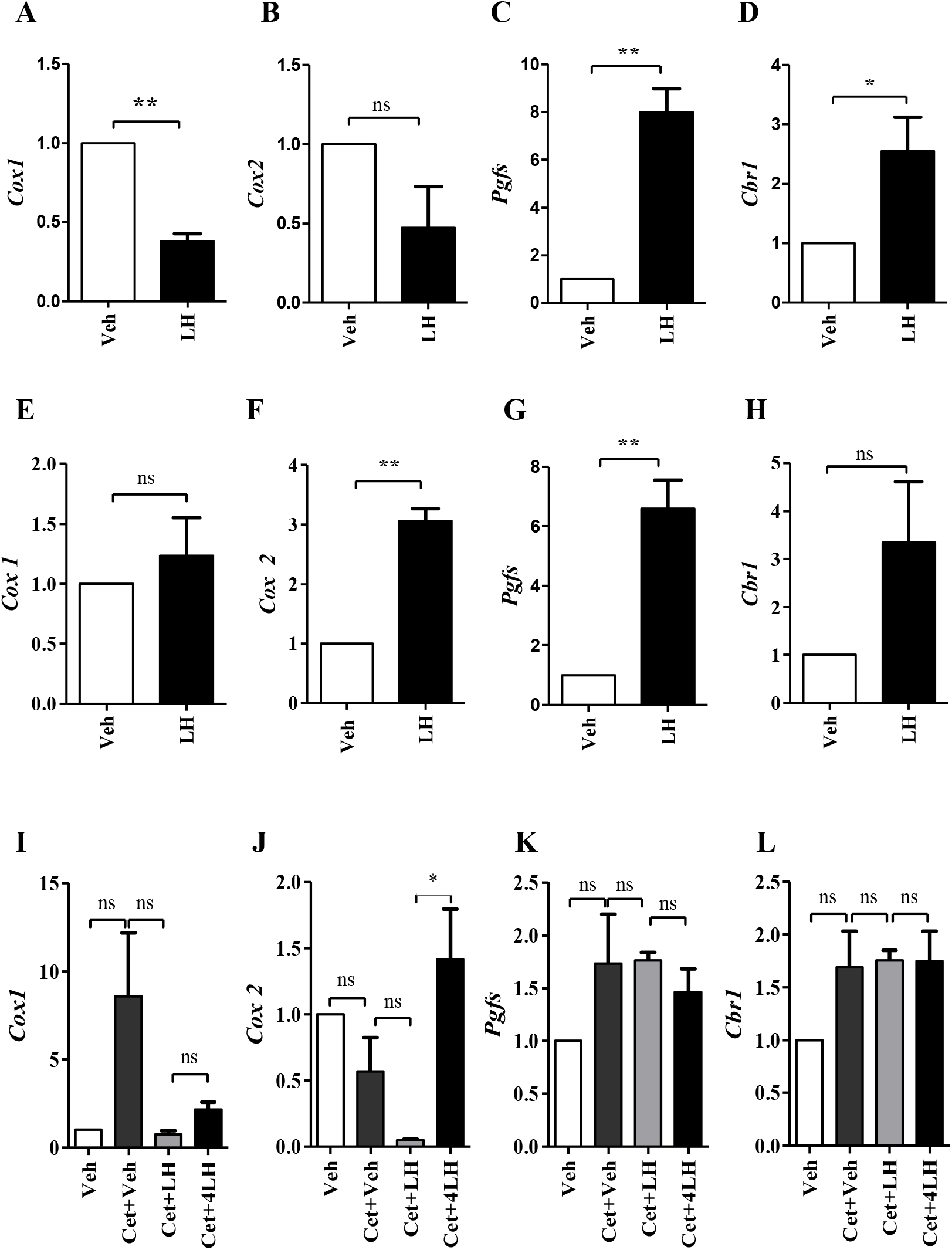
Effects of multiple injections of exogenous LH on prostaglandin synthesis machinery in late and mid-pregnant rats. Luteal *Cox1, Cox2, Pgfs, Cbr1* mRNA expression during late (A-D) and mid-pregnancy (I-L). Uterine *Cox1, Cox2, Pgfs, Cbr1* mRNA expression during late-pregnancy (E-H). L19 mRNA expression levels were used as internal control for qPCR analyses. The data represent mean±S.E.M of 3 rats/group. *<0.05; **<0.01 & ***<0.001.

### Effect of repeated LH administration on genes associated with activated PGF_2α_ signalling and uterine activation during pregnancy

During late-pregnancy, repeated administration of LH significantly decreased luteal expression of *11β-hsd2* (p<0.001) and *Timp3* (p<0.05) compared to Veh treated rats (Fig. 2A, C). However, expression of *Scp2* decreased non-significantly upon repeated LH treatment (Fig. 2B). We analyzed the transcript levels of CAPs and genes involved in uterine relaxation to assess uterine activation. During late-pregnancy, expression of *Il-1β* (non-significant), *Infr-γ* (p<0.01), and *Igfbp2* (p<0.01) increased in the uterine tissue of rats receiving repeated LH administration (Fig. 2D-F). Expression of *Ugn* (p□0.01), *Ep2* (non-significant), and *Igfbp5* (p<0.05) decreased in the uterine tissue post repeated LH administration during late-pregnancy (Fig. G-I). No change occurred in the levels of PGF_2α_ signaling genes in the CL and uterine tissue collected from mid-pregnant rats (data not shown). Expression analysis of PGF_2α_ biosynthesis and its downstream signaling genes suggested that PGF_2α_ signaling is activated in LH induced luteolysis during the late-stage and not during the mid-stage of pregnancy. Therefore, to further examine the role of PGF_2α_ signaling in LH-induced luteolysis, we administered DIC (an inhibitor of PG synthesis) to block the endogenous synthesis of PG during the late-stage of pregnancy in rats.

**Figure 2.**
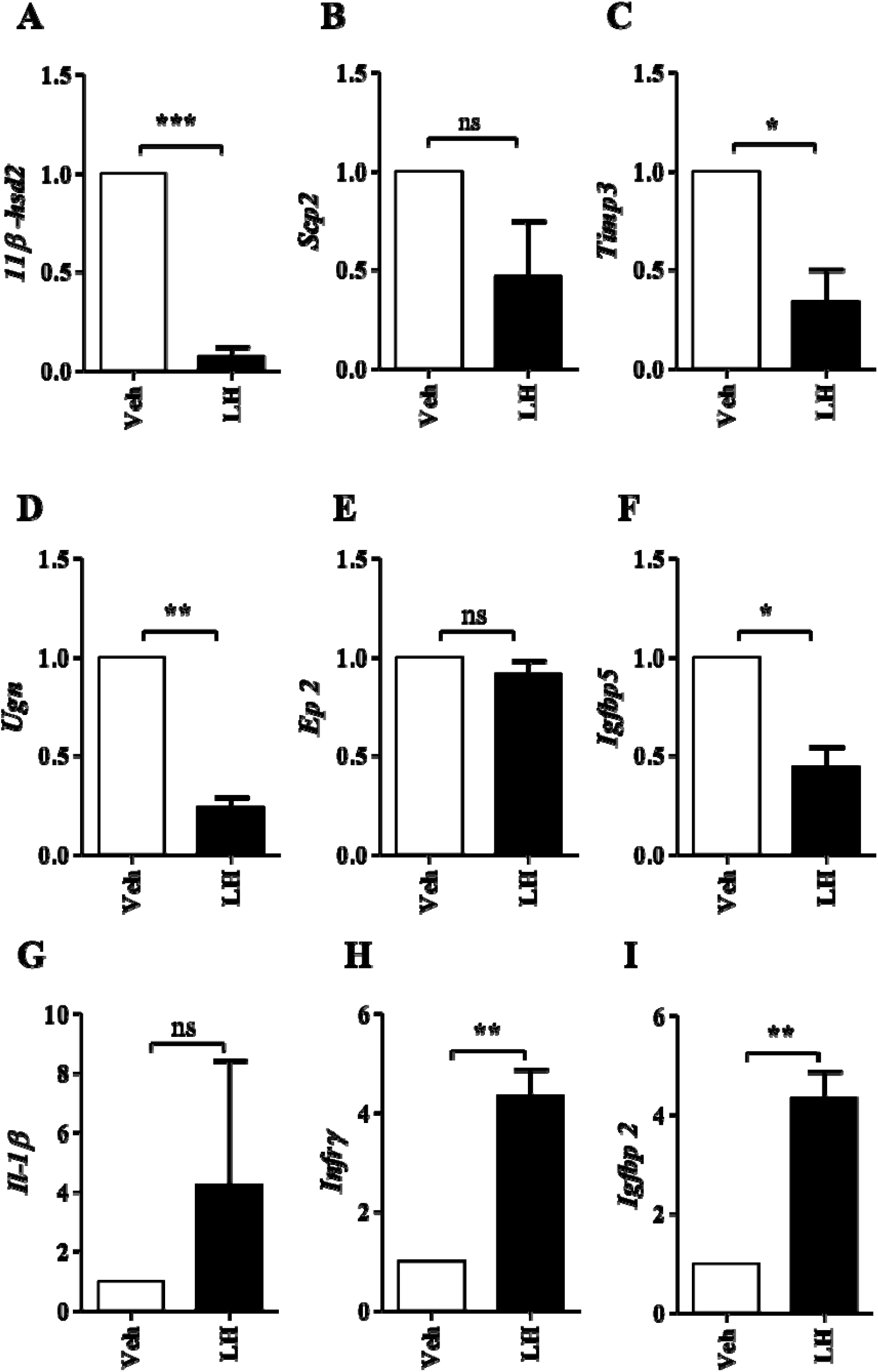
Effects of multiple injections of exogenous LH on genes associated with PGF_2α_ signalling during late-pregnancy. Luteal *11β-hsd2* (A), *Scp2* (B), *Timp3* (C) mRNA expression. Uterine *Ugn* (D), *Ep2* (E), *Igfbp5* (F), *Il-1β* (G), *Infr-γ* (H) and *Igfbp2* (I) mRNA expression. L19 mRNA expression levels were used as internal control for qPCR analyses. The data represent mean±S.E.M of 3rats/group. *<0.05; **<0.01 & ***<0.001.

### Effect of DIC treatment on expression of genes associated with PGF_2α_ biosynthesis during late-pregnancy

DIC treatment reduced luteal expression of genes involved in PGF_2α_ synthesis [*Cox1* (p<0.01), *Cox2* (p<0.01), *Pgfs* (p<0.05), *and Cbr1* (p<0.01)] compared to Veh treated rats (Fig. 3A-D). Expression of *Cox1* (p<0.05), *Cox2* (p<0.01) and *Pgfs* (p<0.01) decreased in the uterus post DIC treatment (Fig. 3E-G). However, expression of *Cbr1* showed non-significant decrease (Fig. 3H). After confirming inhibition of the genes associated with PGF_2α_ biosynthesis by DIC treatment, LH injections were administered to DIC treated rats to examine if LH could still mediate its luteolytic effects in the absence of endogenous PG signaling. The experimental regimen followed is described in the material and methods section. For simplicity, group of rats that received only DIC treatment will be referred to as DIC group and the group that received a combination of DIC plus 4 injections of LH will be referred to as DIC+4LH group henceforth.

**Figure 3.**
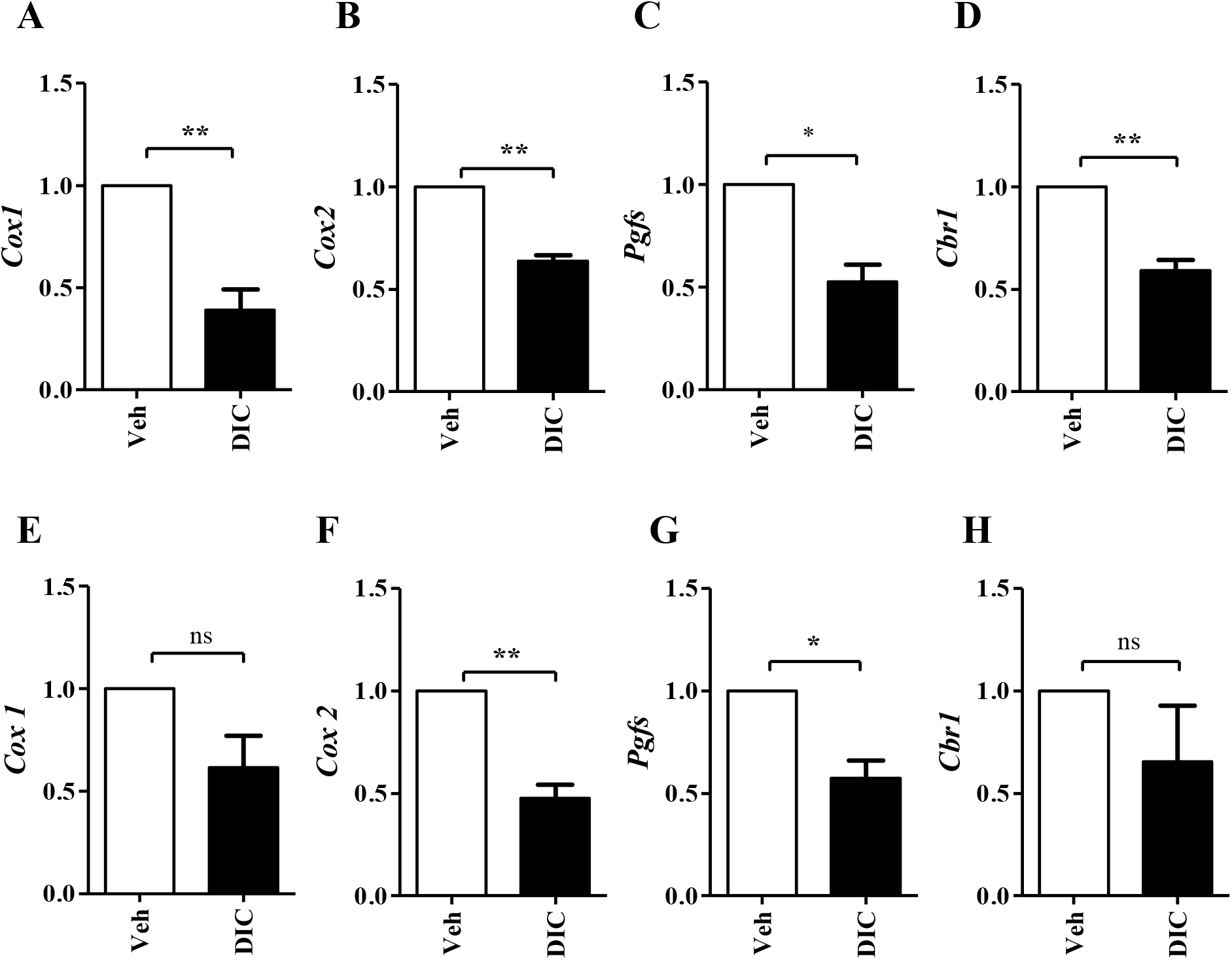
Effects of DIC treatment on prostaglandin synthesis machinery during late-pregnancy. Luteal *Cox1* (A), *Cox2* (B), *Pgfs* (C), *Cbr1* (D) mRNA expression. Uterine *Cox1* (E), *Cox2* (F), *Pgfs* (G), *Cbr1* (H) mRNA expression. L19 mRNA expression levels were used as internal control for qPCR analyses. The data represent mean±S.E.M of 3 rats/group. *<0.05; **<0.01 & ***<0.001.

### Effect of repeated LH treatment on expression of PGF_2α_ biosynthesis genes uterine activation in DIC pre-treated late-pregnant rats

Luteal expression of *Cox2* (p<0.01), *Pgfs* (p<0.05) and Cbr*1* (non-significant) increased in the DIC+4LH compared to DIC group (Fig. 4B-C). However, the expression of *Cox1*, the constitutively expressed isoform, decreased in the DIC+4LH group compared to DIC group (Fig. 4A). The expression of *Cox1* (non-significant), *Cox2* (non-significant), *Pgfs* (p<0.01) and *Cbr1* (non-significant) increased in the uterine tissue collected from DIC+4LH group (Fig. 4E-H). The expression of *CAPs [Il1-β* (p<0.01), *Infr-γ* (p<0.05) and *Igfbp2* (non-significant)] increased in the DIC+4LH group compared to DIC group (Fig 5A-C). The expression of genes associated with uterine relaxation [*Ugn* (p<0.001), *Ep2* (p<0.05) and *Igfbp5* (non-significant)] decreased in the uterine tissue collected from the DIC+4LH group compared to DIC group (Fig. 4D-F).

**Figure 4.**
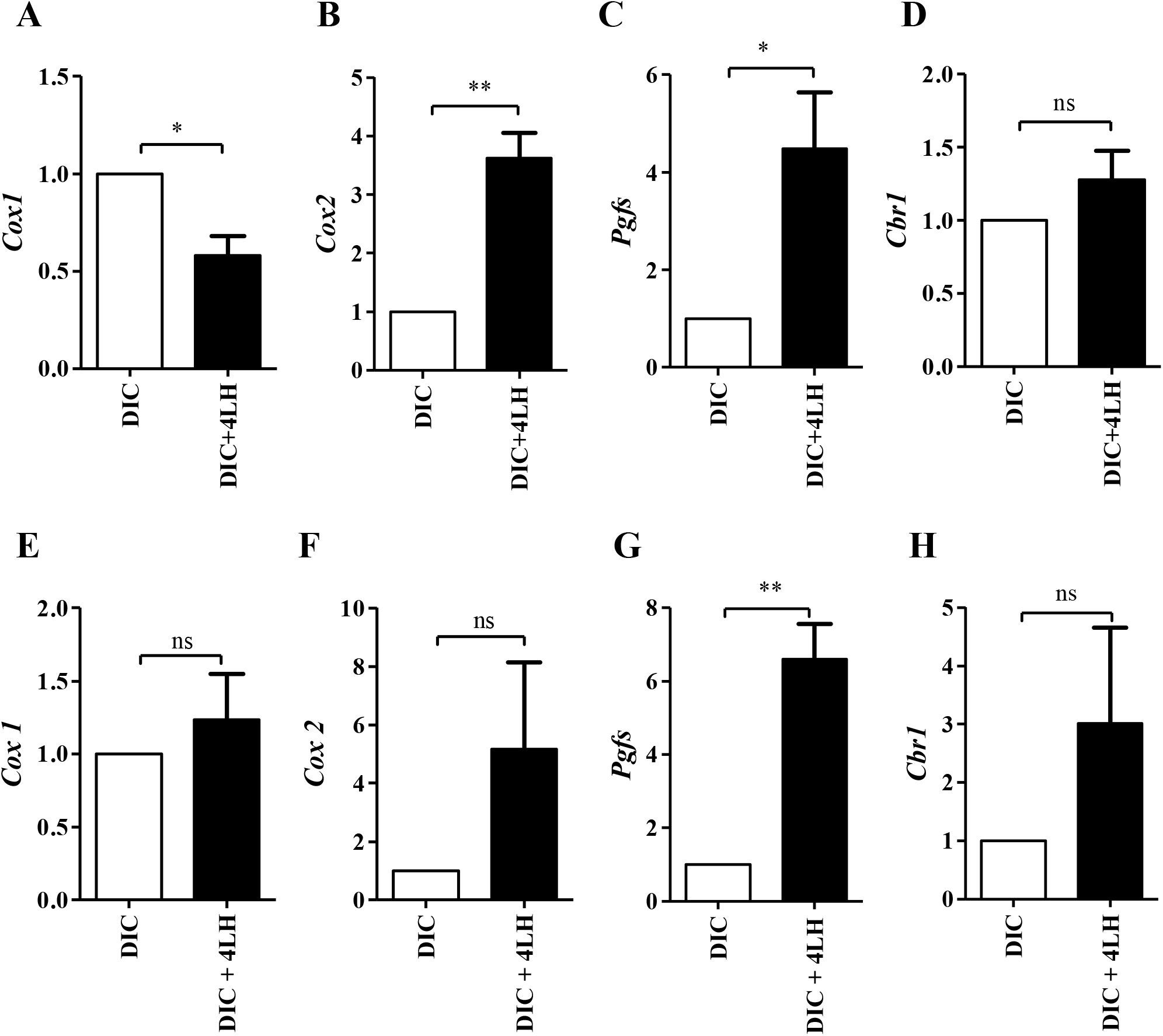
Effects of multiple injections of exogenous LH on prostaglandin synthesis machinery during late-pregnancy. The rats were also treated with DIC to inhibit endogenous prostaglandin secretion prior to administration of single or multiple injections of LH. Luteal *Cox1* (A), *Cox2* (B), *Pgfs* (C), *Cbr11* (D) mRNA expression. Uterine *Cox1* (E), *Cox2* (F), *Pgfs* (G), *Cbr1* (H) mRNA expression. L19 mRNA expression levels were used as internal control for qPCR analyses. The data represent mean±S.E.M of 3 rats/group. *<0.05; **<0.01 & ***<0.001.

**Figure 5.**
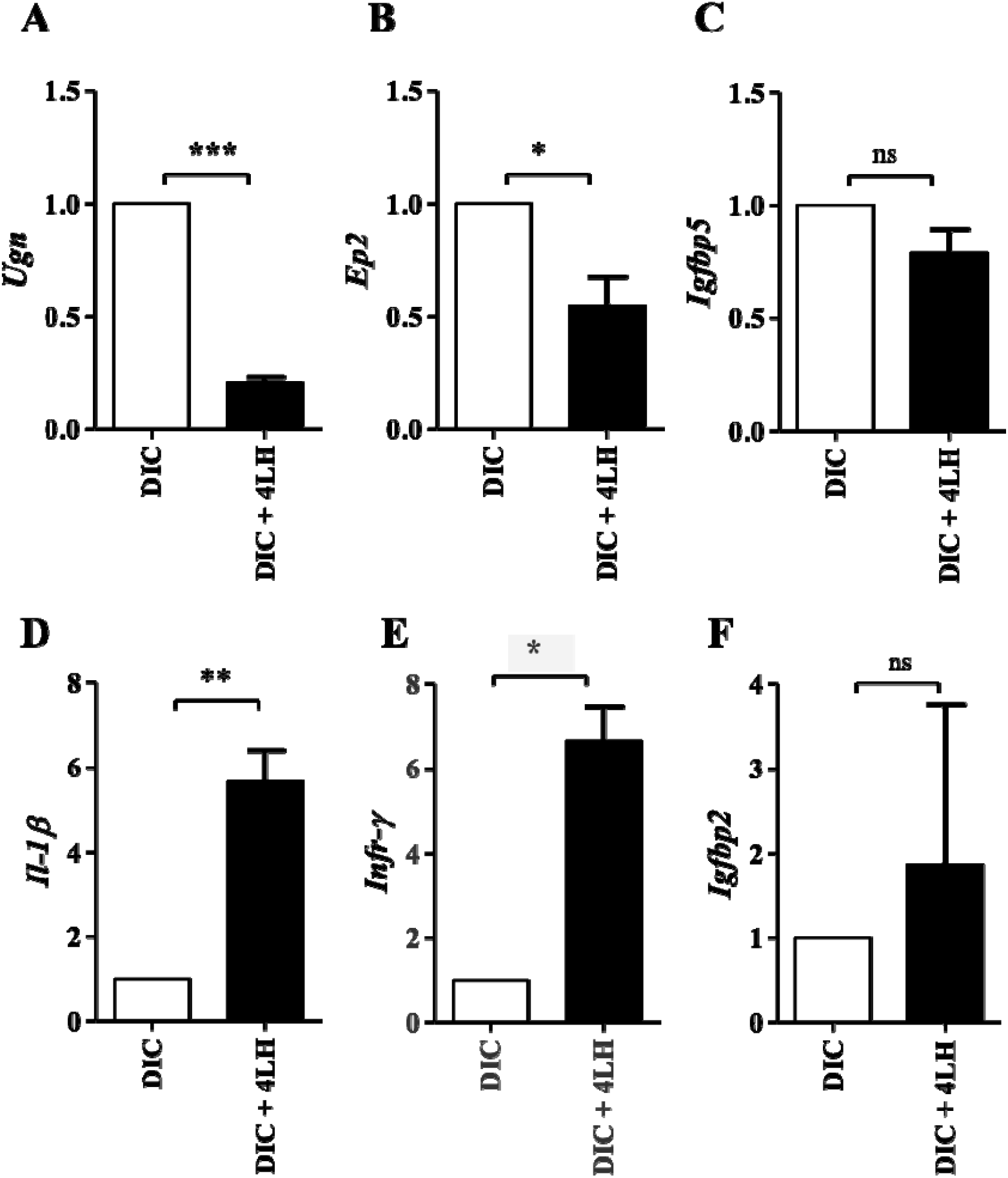
Effects of multiple injections of exogenous LH on genes associated with PGF_2α_ signalling during late-pregnancy. The rats were also treated with DIC to inhibit endogenous prostaglandin secretion prior to administration of single or multiple injections of LH. Uterine *Ugn* (A), *EP2* (B), Igfbp5 (C), *Il-1β* (D), *Infr-γ* (E) and *Igfbp2* (F) mRNA expression. L19 mRNA expression levels were used as internal control for qPCR analyses. The data represent mean±S.E.M of 3rats/group. *<0.05; **<0.01 & ***<0.001.

### Effect of repeated LH treatment on cAMP/PKA/CREB pathway in CL of rats treated with DIC

The down-regulation of cAMP/PKA/CREB pathway post repeated LH treatment plays a crucial role in LH-mediated luteolysis (Vashistha *et al*., 2021). We next analysed the effect of blockage of endogenous PGs on LH induced down regulation of cAMP/PKA/CREB pathway. As can be seen in Fig. 6A-C, repeated administration of LH to late-pregnant rats decreased the luteal expression of Lh/cgr (p<0.01), cAMP content (p<0.01) and pCREB protein levels (p<0.01) compared to Veh-treated rats. Inhibition of endogenous PG biosynthesis (DIC group) did not affect the cAMP/PKA/CREB pathway as revealed by no significant change in Lh/cghr transcript levels, luteal cAMP levels and phosphorylation status of CREB in comparison to Veh treated animals (Fig. 6A-C). We observed a reduction in *Lh/cghr* (p<0.001) expression, cAMP levels (p<0.05) and pCREB (non-significant) in the luteal tissue of DIC+4LH treated rats compared to DIC treated group (Fig. 6A-C). However, rats that received either LH or a combination of DIC and LH (DIC+4LH group) had no significant difference in the mRNA levels of *Lh/cgr*, luteal cAMP content and phosphorylated CREB (Fig. 6A-C). The results indicate that repeated LH treatment following inhibition of endogenous PGF_2α_ synthesis can still causes down-regulation of the cAMP/PKA/CREB pathway. This suggests that endogenous PGF_2α_ does not play a role in LH-induced down-regulation of the cAMP/PKA/CREB pathway.

**Figure 6.**
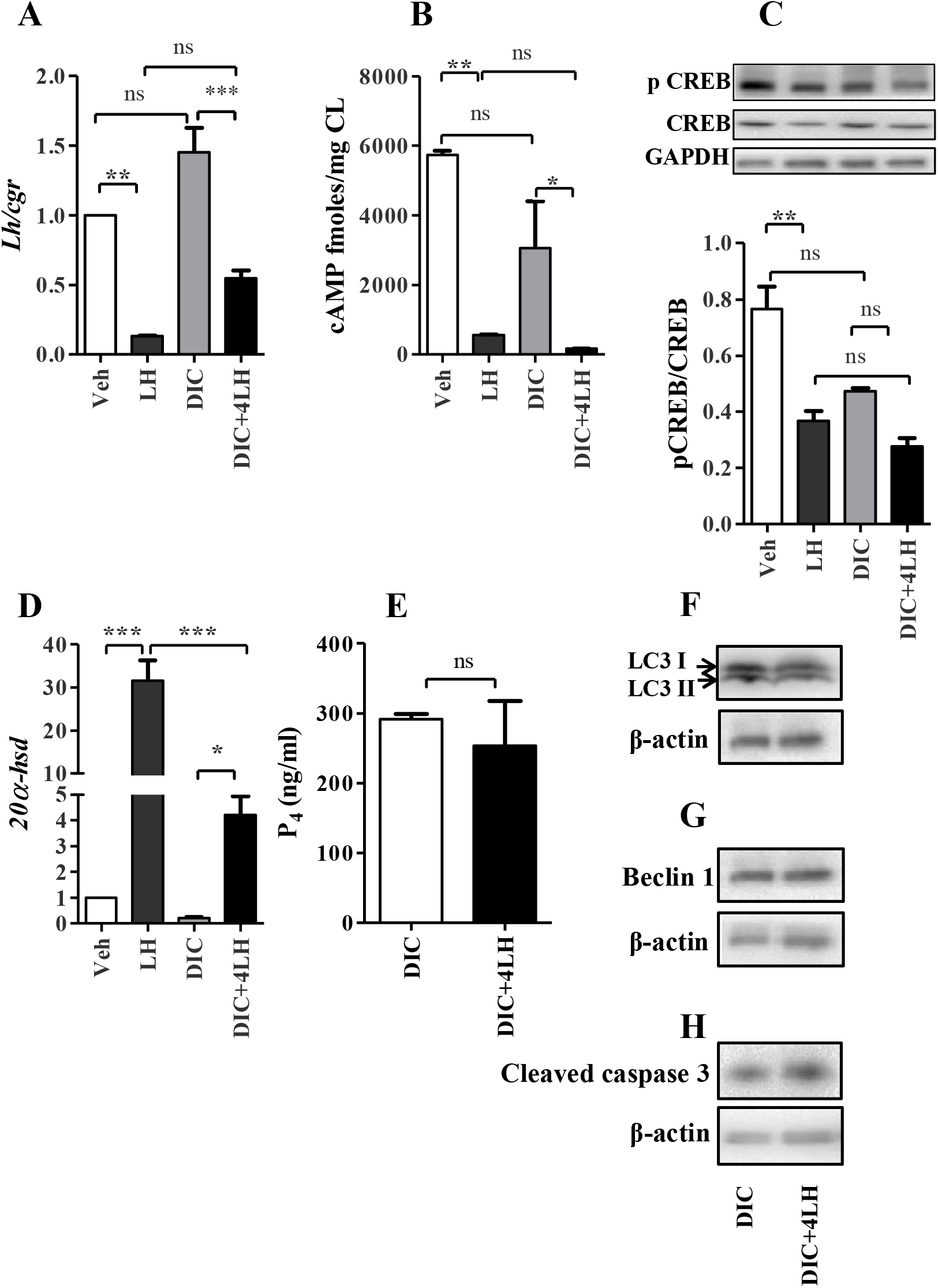
Effects of multiple injections of exogenous LH on various components of cAMP signal transduction pathway and markers of luteolysis in late-pregnant rats. The rats were also treated with DIC to inhibit endogenous prostaglandin secretion prior to administration of single or multiple injections of LH. mRNA expression of *Lh/cgr* (A), cAMP levels (B), representative immunoblot analysis image and quantitation data for pCREB and CREB (C), mRNA expression of luteal *20*α*-hsd* (D) and P_4_ serum levels (E), representative immunoblot analysis image and quantitation data for LC3 II (F), Beclin 1 (G) and Cleaved Cas 3 (H). L19 mRNA expression was used as an internal control for qPCR. GAPDH and β-actin protein levels were used as internal controls for qPCR and immunoblot analyses. Data represented mean ± S.E.M of 3-5 rats/group. *< 0.05; ** <0.01 & *** < 0.001.

### Effect of repeated LH treatment on functional and structural luteolysis in rats treated with DIC

Analysis of the effect of inhibition of endogenous PG biosynthesis on LH-mediated functional luteolysis revealed that the expression of the luteolytic marker *20*□*-hsd* increased (p<0.001) upon repeated administration of LH compared to Veh treated group (Fig. 6D). Inhibition of endogenous prostaglandins did not significantly alter its expression compared to the Veh treated group (Fig. 6D). An increase (p<0.05) in its expression was observed in the DIC+4LH group compared to the DIC group (Fig. 6D). However, the increase in the expression of the gene *20*□*-hsd* in the DIC+4LH group was significantly less than the group in which endogenous PGs were not depleted (LH injections group) (Fig. 6D). Nonetheless, the findings indicate that endogenous PG play some role in functional luteolysis initiated by repeated LH treatment during the late-stage of pregnancy. However, we did not observe any change in the serum P_4_ levels between the DIC and DIC+4LH group (Fig. 6E). Further analysis of the effect of depletion of endogenous PGF_2α_ on LH-mediated structural luteolysis was carried out by analyzing the protein levels of apoptosis (Cleaved caspase 3) and autophagy (LC3I/II, Beclin1) markers. Protein levels LC3I/II, Beclin1 and Cleaved caspase 3 did not change in the luteal tissue collected from the DIC+4LH group compared to DIC group (Fig. 6F-H). The findings suggest that repeated LH administration treatment protocol in the absence of endogenous PGF_2α_ signaling activates only functional luteolysis.

## Discussion

Repeated administration of LH initiates the process of luteolysis during different stages of pregnancy in rats (Vashistha *et al*., 2021). Our study explains the effect of repeated LH administration on luteal and uterine PGF_2α_ biosynthesis machinery, uterine activation and the effect of blockage of overall PG synthesis machinery on LH-mediated luteolysis.

LH is known to have stage dependent effects on pregnancy (Vashistha *et al*., 2021). We did not observe any significant change in the expression of PGF_2α_ synthesis and PGF_2α_ responsive genes in the luteal and uterine tissue post repeated LH treatment during the mid-pregnancy. The results suggests that endogenous PGF_2α_ signaling may not play a significant role in the LH induced luteolysis during mid-pregnancy (Farina *et al*., 2004). It was demonstrated in the lab previously that the serum levels of P_4_ do not change upon repeated LH treatment during mid-stage of pregnancy (Vashistha *et al*., 2021). The limited role of PGF_2α_ signaling during mid-pregnancy can be explained as the inhibitory effects of P_4_ are still intact and prevents the activation of PGF_2α_ signaling during repeated LH treatment. P_4_ has been demonstrated to play a crucial role in maintaining uterine quiescence by various groups (Farina *et al*., 2004). In several mammalian species, excepting humans, a drop in P_4_ is a prerequisite for the onset of labor (Kota *et al*., 2013). Various studies corroborate that a drop in P_4_ levels is required to initiate the luteolytic events and uterine activation (Stocco and Deis, 1996; Kota *et al*., 2013). P_4_ has been shown to mediate its effect of maintaining uterine quiescence through a number of mechanisms which include increase in the expression of calcium and potassium channels (Mendelson and Condon, 2005; Soloff *et al*., 2011), down regulation of expression of uterotonic agents and members of their signaling cascades (Mendelson and Condon, 2005; Soloff *et al*., 2011) and decrease in the cross-linking of the actin with the myosin filament (Soloff *et al*., 2011).

In the present study, repeated LH stimulation decreased the expression of *Cox1* and *Cox2* in the late-stage pregnant rat luteal tissue suggesting a downregulation in overall prostaglandin production. However, an increase in the expression of *Pgfs* and *Cbr1* suggests that repeated LH administration specifically shifts the prostaglandin synthesis machinery towards PGF_2α_ production. Other effectors of PGF_2α_ signaling that are known to regulate steroidogenesis include *11β-hsd2, Scp2* and *Timp3* (Berridge and Irvine, 1984). We observed a decrease in their expression upon repeated LH treatment. These genes get down-regulated on PGF_2α_ treatment (Berridge and Irvine, 1984; Hardy *et al*., 1999) and hence serve as the markers of increased PGF_2α_ signaling. Unlike the mid-stage, the results obtained during the late-stage suggest that the repeated administration of LH stimulates PGF_2α_ signalling in the CL. Similar observations were made by Stocco et.al., where they demonstrated an increase in luteal prostaglandin content which further led to an increase in 20α-HSD activity in LH-induced luteolysis during the late-stage of pregnancy in rats (Stocco and Deis, 1998).

In rats, luteolysis precedes the process of parturition. For the initiation of parturition, the uterus transforms from a quiescent state to a highly active state. This transformation involves increased expression of genes encoding prostaglandin synthesis machinery, CAPs and repression of uterine relaxant systems (JRG *et al*., 2000; Mitchell *et al*., 2005). During the late-stage of pregnancy, an increase in the expression of uterine prostaglandin synthesis machinery was observed post repeated LH administrations. Similar observations have been made by others as well in leydig cells (Chen *et al*., 2007). In bovines, LH stimulates the expression of *Cox2, Pgfs* and *Cbr1* (Chen *et al*., 2007).

In the present study, the expression of CAPs increased, and the expression of genes involved in uterine relaxation decreased post LH administration during the late-stage of pregnancy. The results suggest that LH-mediated luteolysis activates PGF_2α_ signaling in the uterine tissue. An increase in the prostaglandin signaling during luteolysis and parturitions has been observed by others as well. Among the CAPs, *Ugn* is known to undergo rapid reduction just prior to parturition (Girotti and Zingg, 2003). EP receptor subtypes EP1, EP2, EP3 and EP4 receptors have different effects on the uterus: EP1 and EP3 are involved in uterine contraction while EP2 and EP4 are responsible for uterine relaxation (Astle *et al*., 2005). We observed a decrease in the expression of EP2 receptor subtype suggesting activation of the uterus. Brodt-Eppley and colleagues (Astle *et al*., 2005) analysed the expression of EP2 receptor throughout gestation and observed that the mRNA levels of EP2 were highest on day 16 of pregnancy and declined significantly until parturition. We observed an increase in the levels of cytokines post repeated LH administration. An increase in the levels of cytokines during PGF_2α_ induced luteolysis in bovine corpus luteum has been reported by others (Chen *et al*., 2007).

Repeated LH administration has been shown to activate luteolytic events both *in-vivo* and *in-vitro* via desensitization of the cAMP/PKA/CREB pathway (Vashistha *et al*., 2021). PGF_2α_ is a well-established luteolysin in rodents and is known to interact with the cAMP/PKA/CREB pathway (Astle *et al*., 2005). Hence, we analysed the effect of inhibition of endogenous PGs on the cAMP/PKA/CREB pathway. Inhibition of endogenous PG synthesis did not have any effect on cAMP/PKA/CREB pathway. Similar effects on cAMP/PKA/CREB pathway upon stimulation with PGF_2α_ have also been reported in bovine luteal cell culture (Astle *et al*., 2005). The enzyme AC generally acts as the site for interaction between the prostaglandin signaling and cAMP/PKA/CREB pathway (Astle *et al*., 2005). Out of the 9 ACs, AC 1, AC 3 and AC 8 are known to be stimulated by Ca^2+^ released on FP receptor activation. AC 5, AC 6 are inhibited whereas AC 2, AC 4, AC 7 and AC 9 are unaffected by Ca^2+^ (Astle *et al*., 2005). This suggests that the type of interaction between the FP receptor and adenylate cyclase pathway might depend on the AC isoform expressed.

PGF_2α_ positively regulates the expression of the luteolytic marker *20*α*-hsd* (Stocco *et al*., 2000). In rats, PGF_2α_ administration induces premature expression of 20α-HSD in the luteal tissue (Stocco *et al*., 2000). The increased expression of 20α-HSD gives the signal for parturition in rodents. The primary function of 20α-HSD is to increase the catabolism of P_4_ into an inactive metabolite 20α-DHP (Stocco *et al*., 2000). Our results indicate that even in the absence of endogenous PGF_2α_ signaling, repeated LH treatment could initiate functional luteolysis but not structural luteolysis. This suggests that desensitization of the cAMP pathway and an active PGF_2α_ signaling are required for the initiation of luteolysis to full extent.

In summary, our results suggest that endogenous PGF_2α_ may contribute to LH-mediated luteolysis, but this dependency on endogenous PGF_2α_ is pregnancy stage dependent. Unlike the mid-stage of pregnancy, endogenous PGF_2α_ plays some role during the late stage. Even during the late stage of pregnancy, the participation of PGF_2α_ signaling appears to be subtle compared to the cAMP/PKA/CREB pathway in the initiation of LH-mediated luteolysis.

